# Sex chromosome evolution via two genes

**DOI:** 10.1101/494112

**Authors:** Alex Harkess, Kun Huang, Ron van der Hulst, Bart Tissen, Jeffrey L. Caplan, Aakash Koppula, Mona Batish, Blake C. Meyers, Jim Leebens-Mack

**Affiliations:** Donald Danforth Plant Science Center, St. Louis MO 63132; Department of Plant Biology, University of Georgia, Athens GA; Delaware Biotechnology Institute, University of Delaware, Newark, DE 19716; Limgroup B.V., Veld-Oostenrijk 13, 5961 NV Horst, Netherlands; Department of Biological Sciences, University of Delaware, Newark, DE 19716; Department of Medical and Molecular Sciences, University of Delaware, Newark DE 19716; Division of Plant Sciences, University of Missouri, Columbia MO 65211

## Abstract

The origin of sex chromosomes has been hypothesized to involve the linkage of factors with antagonistic effects on male and female function. Garden asparagus (*Asparagus officinalis* L.) is an ideal species to test this hypothesis, as the X and Y chromosomes are cytologically homomorphic and recently evolved from an ancestral autosome pair in association with a shift from hermaphroditism to dioecy. Mutagenesis screens paired with singlemolecule fluorescence *in situ* hybridization (smFISH) directly implicate Y-specific genes that respectively suppress female organ development and are necessary for male gametophyte development. Comparison of contiguous X and Y chromosome shows that loss of recombination between the genes suppressing female function (*SUPPRESSOR OF FEMALE FUNCTION, SOFF*) and promoting male function (*TAPETAL DEVELOPMENT AND FUNCTION 1, aspTDF1*) is due to hemizygosity. We also experimentally demonstrate the function of *aspTDF1*. These finding provide direct evidence that sex chromosomes can evolve from autosomes via two sex determination genes: a dominant suppressor of femaleness and a promoter of maleness.

**One Sentence Summary:** Sex chromosomes can evolve from autosomes by mutations in two sex determination genes.

## MAIN TEXT

Beginning with Nettie Stevens’ elegant explanation of male and female mealworm gamete differences involving “sex chromosomes” in 1905 (*1*), sex chromosomes have been characterized in dioecious species (with separate male and female sexes) across all major eukaryotic lineages. Flowering plants offer a unique perspective to the origin and early evolution of sex chromosomes across the tree of life, given that dioecy has evolved hundreds of independent times from hermaphroditic ancestors across the angiosperm phylogeny (*2*).

Several models have been proposed to explain the transition from a hermaphroditic species with autosomal chromosomes into a dioecious species in association with the evolution of an X/Y or Z/W sex chromosome pair. One model developed over time by both Westergaard (*3*, *4*) and Charlesworth & Charlesworth (*5*) hypothesizes that a sex chromosome might be able to evolve from an autosomal pair via mutations in two perfectly linked genes in a non-recombining region of a proto-Y chromosome: one gene that promotes anther development, and another that suppresses female organ development. This model has been notoriously difficult to test in plants and animals, given the difficulty of assembling sex chromosomes and identifying causal sex determination genes, particularly in older sex chromosomes that have degenerated, accumulated repetitive DNA and heterochromatin, undergone structural variations, and possibly experienced gene turnovers. Here we show functional evidence that in garden asparagus (*Asparagus officinalis* L.), the recently evolved Y chromosome harbors two sex determination genes that control flower sex expression.

## RESULTS AND DISCUSSION

### Two Y-linked genes determine sex in garden asparagus

Sequencing of a YY “supermale” garden asparagus genome revealed existence of a nearly one megabase non-recombining, male specific sex determination region (SDR) on the Y chromosome with only thirteen gene models (*6*), two of which we identify here as the sex determination genes (Figure 1A). Previous work to identify the sex determination genes in the non-recombining regions of the garden asparagus X and Y chromosomes utilized gamma irradiation and spontaneous mutants to identify a Y-specific, dominant female suppression gene specifically (*6*). In that experiment, gamma irradiation knockouts of the entire ~1 megabase Y-linked sex determination region (Y-SDR) resulted in male-to-female conversion, whereas single gene knockouts of the Y-specific *SUPPRESSOR OF FEMALE FUNCTION* (*SOFF*) gene converted XY males to hermaphrodites with functioning styles and receptive stigmas (Figure 1B). A single copy gene responsible for proper tapetum development in *Arabidopsis thaliana* (*TAPETAL DEVELOPMENT AND FUNCTION 1, TDF1*) has a male-sterile knockout phenotype (*7*), exists in the non-recombining region, is male-specific across several dioecious *Asparagus* species (*8, 9*), and thus is a strong candidate for a master regulator of male sex determination. However, there was no prior functional evidence in asparagus to support this hypothesis.

Here we use ethyl methanesulfonate (EMS) mutagenesis of an all-XY male population, a male-to-neuter individual that lacked anthers was identified (Figure 1B; Supplemental Figure 1). Resequencing of the Y-specific *aspTDF1* gene in this mutagenized individual revealed a single nucleotide polymorphism that induced a premature stop codon in the predicted protein (Supplemental Figure 1). Functional modification of one or both of the sex-determining *aspTDF1* and *SOFF* genes can convert XY males into three different sexual forms: knockout of the *SOFF* gene converts males to hermaphrodites, knockout of the *aspTDF1* gene converts males to neuters, and knockout of both *TDF1* and *SOFF* converts males to females (Figure 1B). With knockouts for two sexually antagonistic Y-linked genes, one that promotes anther formation (*TDF1*) and another that suppresses pistil development (*SOFF*), this is the first functional evidence in a sex chromosome-bearing dioecious species supporting the long-standing “two gene” model of sex chromosome evolution from Westergaard (*3*) and Charlesworth and Charlesworth (*10*).

**Fig. 1.**
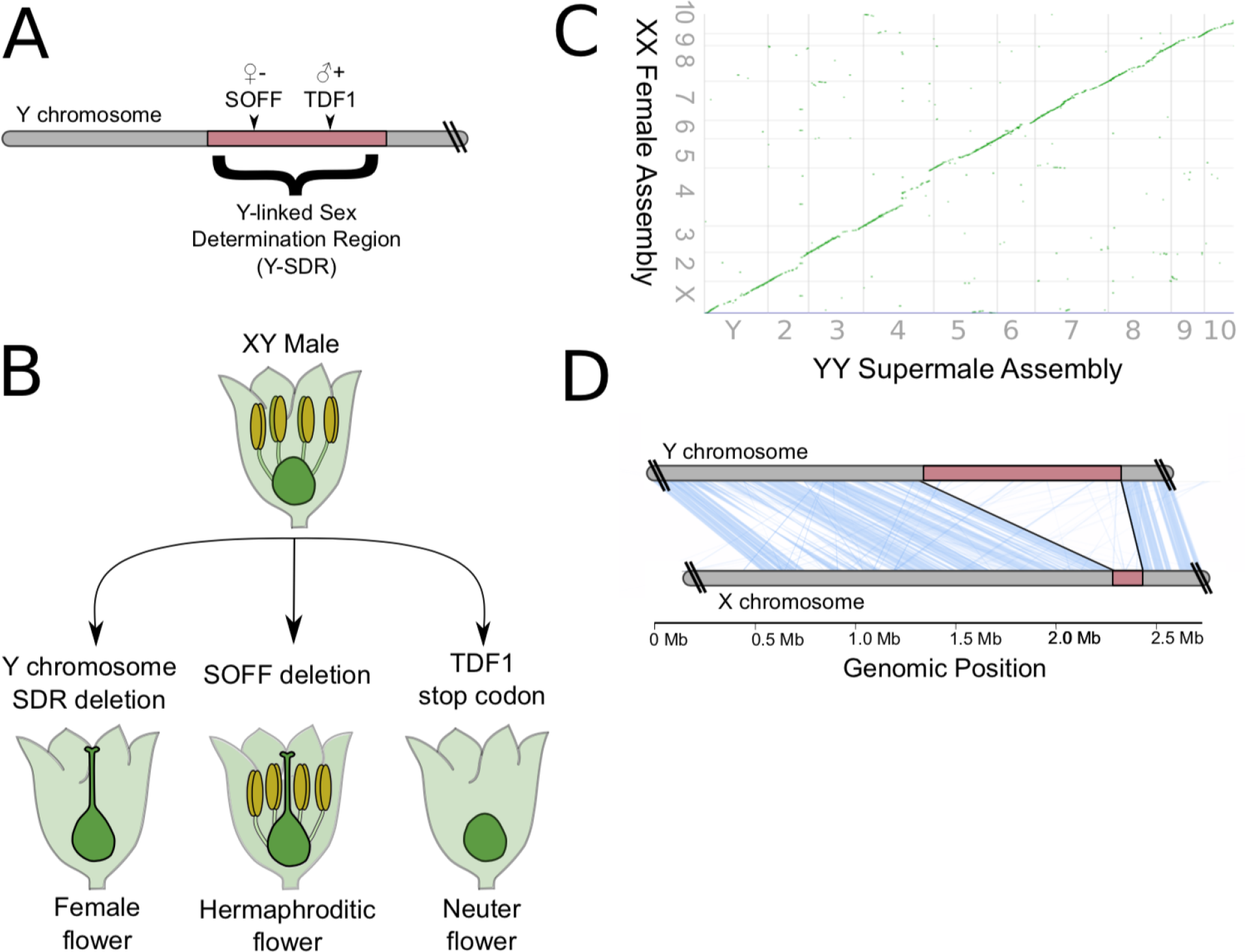
The structure and sex-determining function of the garden asparagus Y chromosome and its structural variation relative to the X chromosome. (A) Depiction of the Y chromosome telomeric region and the nearly 1 megabase Sex Determination Region (SDR). Two Y-linked sex determining genes, *SUPPRESSOR OF FEMALE FUNCTION* (*SOFF*) and *DEFECTIVE IN TAPETAL DEVELOPMENT AND FUNCTION 1* (asp *TDF1*) are contained in this non-recombining region. SOFF dominantly suppresses female organogenesis, whereas TDF1 promotes proper anther development. Both genes are missing from the X chromosome. (B) Sexual conversions are possible by functionally modifying the Y-SDR. Gamma irradiations that delete the entire Y-SDR region convert XY males into females. Single-gene gamma irradiation mutants of *SOFF* convert XY males into hermaphrodites. A premature stop codon in asp*TDF1* converts XY males into neuters. (C) Synteny of the XX PacBio and BioNano genome assembly against the previous YY genome assembly across all 20 chromosomes (*6*). (D) Microsynteny of the X chromosome against the Y chromosome (blue lines) with the non-recombining regions of both chromosomes (red blocks). Blue lines indicate nucmer alignment matches greater than 1.5 kb with 90% minimum identities.

### The structure of the X chromosome

One necessary feature of plant sex chromosomes is a mechanism to suppress recombination between the heterologous pair, maintaining the sex determination genes in perfect linkage. Given the multiple independent origins of dioecy and sex chromosomes across flowering plants (*2*), it is unsurprising that suppressed recombination between X/Y or Z/W can initiate and persist via multiple mechanisms. For example, in papaya (*Carica papaya*), the non-recombining region of the Y has repeatedly inverted and expanded across the already non-recombining centromere (*11*). In poplar trees (*Populus sp.*), a translocation event of a sex-specific region to the distal tip of chromosome 19 inhibits recombination between the sex chromosome pair (*12, 13*). These initial events may be followed by extensive Y-linked repeat proliferation, exemplified by the large and expanded heterochromatic Y chromosomes in *Silene latifolia* and *Coccinia grandifolia* (*14*).

Previous inferences about the structure and composition of the garden asparagus X chromosome have relied on resequencing data aligned to the YY reference genome. We generated a *de novo* whole genome assembly of a doubled-haploid XX garden asparagus individual that is a sibling of the previously assembled YY genome. Assembly of roughly 40X coverage PacBio reads with FALCON yielded the expected 1.286 gigabase (Gb) genome assembly with 5,647 contigs (contig N50 = 385 kb), and further scaffolding with a BioNano Genomics BspQI optical map reduced the scaffold count to 4,502 (scaffold N50 =1.67 Mb). The assembly was anchored against the YY genome pseudomolecules (Figure 1B). BUSCO metrics (*15*) show that the XX genome assembly has 85.3% of full-length embryophyte database orthologs, nearly the same as the previous YY assembly (Supplemental Figure 2), suggesting that it is highly complete.

The X chromosome assembly contained a contiguously assembled 163 kb region of X-specific sequence that corresponds with the sex determination region on the Y (Figure 1C). Compared to the nearly 1 Mb Y-specific region, this shows that hemizygosity is the primary driver of non-recombination between X and Y in garden asparagus. The *SOFF* and *aspTDF1* genes are not present on the homologous X region or elsewhere in the XX female genome. There is no evidence of repeated inversions leading to the formation of strata on the asparagus Y chromosome (Figure 1D), as has been inferred for older plant sex chromosomes like *Carica papaya* (*11*) and *Silene latifolia* (*16*, *17*). Repetitive elements on the garden asparagus X chromosome are dominantly Long Terminal Repeat (LTR) retrotransposons with an insertion age distribution similar to that of entire gene (Supplemental Figure 3), suggesting that the non-recombining region of the X chromosome is able to purge LTR elements via recombination with the homologous X.

One possibility during sex chromosome evolution is that genes with male-biased function will accumulate on the Y, and likewise genes with female-biased function will accumulate on the X. Translocations of entire sex-determination regions are even possible (*18*). The 163 kb X-linked region contains five expressed transcripts: three are annotated as uncharacterized proteins, one is a chloroplast-encoded gene shared with the Y, and one is X-linked with no Y homolog (Supplemental Table 1). This annotated X-linked gene is an *Arabidopsis* homolog of NO TRANSMITTING TRACT (WIP2/NTT), a CH2H/C2HC zinc finger transcription factor that in *Arabidopsis thaliana* is specifically expressed in the transmitting tract and funiculus of ovules, playing a key role in the development of fruit (*19*, *20*). Knockouts of WIP2/NTT in *A. thaliana* result in plants that inhibit the movement of the pollen tubes through the carpel and into ovules, severely reducing or eliminating male fertility.

### Single Molecule FISH expression quantification

Previous attempts at RNA-seq of whole spear tips (with both vegetative and floral buds) were able to clearly detect the expression of *aspTDF1*, but not *SOFF* (*21*). Little is known about the mode of action of the SOFF protein, other than it containing a PFAM annotation “Domain of Unknown Function 247 (DUF247)”. Intriguingly, a DUF247 domain-containing protein co-segregates with the S Self-Incompatibility locus in perennial ryegrass (*22*), hinting at the possibility of a conserved potential pistil suppression function. To clarify the spatial expression patterns of both genes to better elucidate their respective functions, single-molecule fluorescence *in situ* hybridization (sm-FISH) was used to track the expression and localization of the sex-determination genes on XX and YY spear tips. Since the two sex determination genes are not present in females, the XX spear tips serve as a negative control. As expected from *Arabidopsis (7), in situs* show that expression of *aspTDF1* is limited to the anther tapetal layer in developing YY supermale buds, and that as expected there is no detectable expression in XX female buds (Figure 2A). The *SOFF* gene is weakly expressed in YY supermale buds, and similarly to asp*TDF1*, is not expressed in XX females (Figure 2A). To resolve the expression levels of *SOFF* and *aspTDF1* in flower buds, the smFISH signals for both *aspTDF1* and *SOFF* were quantified through a Z-stack of entire developing flower buds. We were able to detect the average expression of 288 copies of *aspTDF1* and 99 copies of *SOFF* through a Z-stack of single flower buds, highlighting the efficiency of the *SOFF* gene to inhibit female organogenesis even in low mRNA copy number and little spatiotemporal organization relative to *aspTDF1* (Figure 2B).

**Fig. 2.**
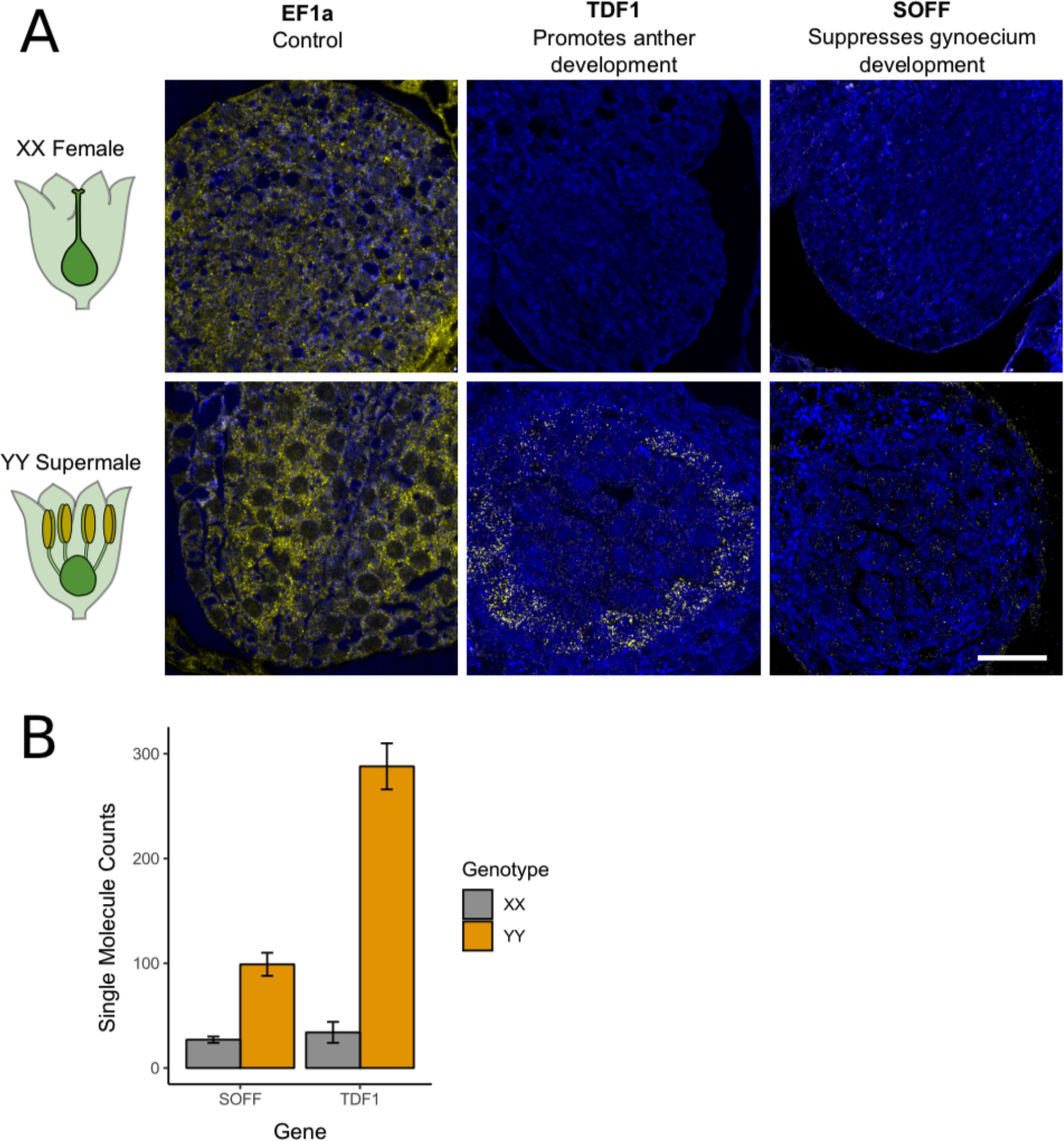
Single Molecule Fluorescence *In Situ* Hybridization (smFISH) of the sex determination genes. (A) smFISH signal for probe (yellow) against a control housekeeping gene EF1a, and the two sex determination genes *aspTDF1* and *SOFF,* in developing pre-meiotic flower buds from spear tips of both an XX female and a YY supermale. Scale bar = 20μm. (B) smFISH molecule expression count for a Z-stack of images through single flower buds for each probe. Errors bars represent standard error.

## Conclusions

Dioecious garden asparagus has a recently evolved sex chromosome that determines sex via the action of two sexually antagonistic genes in perfect linkage on the Y chromosome: a suppressor of female organogenesis (*SOFF*) and a promoter of anther development (*aspTDF1*). Though long hypothesized in several species, this is the first evidence that the evolution of two interacting sex determination genes on a young Y chromosome can evolve to convert a hermaphroditic species into a dioecious one, confirming the long-standing hypothesis posited by Mogens Westergaard (*3*) and Brian and Deborah Charlesworth (*10*).

## Acknowledgements

### Funding

We are thankful for funding from NSF DDIG award #1501589 to JLM and AH, NSF PGRP #1611853 to AH, and NIH #5DP5OD012160-05 to MB.

### Author Contributions

AH, RvdH, BCM, KH and JLM conceptualized the study. AH assembled and annotated the genome. RvdH and BT performed EMS mutagenesis and phenotyping of *tdf1*. KH, JC, MB, and AK performed all imaging and single-molecule FISH analysis.

## Materials and Methods

### EMS mutagenesis

Twelve hundred seeds of the hybrid cultivar Marte (courtesy: Agostino Falavigna) were treated using 0.4 % v/v Ethyl methanesulfonate (EMS). Seeds were imbibed for six hours in rolling flasks filled with 100 mM Na2HPO4. This buffer was replaced by Na2HPO4 buffer plus 0.4% EMS, and seeds were further incubated in rolling flasks for 15 hours at room temperature. EMS treatment was stopped by washing the seeds two times for 20 minutes each in 100 mM Na2S2O3 neutralizing buffer. After neutralization, seeds were rinsed three times in demi water and incubated for another ten minutes in rolling flasks with demi water. Subsequently, the seeds were dried overnight on filter paper in a laminar airflow cabinet at half air speed.

A neuter shoot was discovered on a M1 plant by visual inspection. Meristems of both the WT and mutant shoot taken from the same plant were grown in vitro and rooted as described by Qiao, and Falavigna (1990) to obtain ten cloned plants of each, all of which retained the original phenotype. Sanger sequencing of the TDF1 gene was performed with primer pairs (Forward: AGCCCTTAAGGTTAAATGTCG; Reverse: CATCATGATATAAAATCCTCAATCAAA).

### Asparagus officinalis XX female genome assembly and annotation

High molecular weight DNA was isolated from a doubled haploid XX female sibling to the sequenced YY asparagus genome (*6*) using the Bionano Genomics IrysPrep Plant kit. A 40 kb PacBio library was constructed and size-selected on a SageScience BluePippin, then sequenced with P6C4 chemistry and six hour movies on a Pacific Biosystems RSII. Raw reads greater than 10kb in length were assembled in FALCON. The same high molecular weight DNA was used to generate a BioNano Genomics optical map labeled with Nt.BspQI and imaged to 70X coverage. The assembly was scaffolded against the optical map using the HybridScaffold program in Irysview with default options. The scaffolded assembly was polished using nearly 40X coverage Illumina pair-end 100 nt reads from the same individual with Pilon v1.22, and then assembled into pseudomolecules by comparison to the YY reference genome using CoGE Syntenic Path Assembly. The assembly quality was estimated using BUSCO v3 trained against the maize annotation set.

The assembly was annotated for expressed transcripts using RNA-seq reads from spear tip tissue (which includes developing floral buds) of four XX females (BioProject 259909).

Reads were aligned to the genome with STAR v2.5.3a using default options and --outSAMstrandField intronMotif’, and transcripts were assembled using Stringtie v1.3.3b with default options.

A single X-linked Pacbio contig spanned matched the two pseudoautosomal (PAR) boundaries of the Y. Sequence comparison of the X and Y was performed using MUMMER’s nucmer. The contig was annotated for LTR retrotransposons using LTRharvest, which also computes the relative insertion age of retrotransposons using pairwise LTR end alignments.

### Sample preparation for in situ

Asparagus flower buds of various sizes were cut from the spear and fixed in a 20 ml glass vial using 4% paraformaldehyde in 1x PHEM buffer (5mM HEPES, 60mM PIPES, 10mM EGTA, 2mM MgCl2 pH 7). Fixation was done in a vacuum chamber at 0.08 MPa for 3 times, 15 min each. After fixation, samples were sent for paraffin embedding at the histology lab in the Nemours/Alfred I. duPont Hospital for Children (Wilmington, DE).

### Fluorescence in situ hybridization

Asparagus bud samples were sectioned using a paraffin microtome and dried on poly-l-lysine coated Deckglaser 22×22 mm coverslips (Carl Zeiss Microscopy, LLC, Cat# 474030-9020-000). Samples were then de-paraffinized using Histo-Clear (Fisher Scientific, 50-899-90147) and re-hydrated by going through an ethanol series of 95, 80, 70, 50, 30, 10% (vol/vol) (30 sec each) and water (1 min) at room temperature. After protease (Sigma, P5147) digestion (20 min, 37°C), samples were treated with 0.2% glycine (Sigma-Aldrich, G8898) for 2 min, followed by a TEA (triethanolamine; Sigma-Aldrich, 90279) HCl and acetic anhydride (Sigma-Aldrich, A6404) treatment. After two washes in 1x PBS buffer (phosphate-buffered saline), samples were de-hydrated and then hybridized with smFISH probes. The smFISH probes were designed to bind specifically across the length of the target RNAs (*23*). Briefly, 40 probes (each 20 nt long) were designed for each target, and each probe was synthesized with 3’ amino modification (LGC Biosearch Technologies, CA). All the probes of one set were pooled and *en mass* coupled with TMR (tetramethylrhodamine) or Texas Red and the labeled probe fraction was purified using HPLC (*24*). The probes were diluted in hybridization buffer containing 10% formamide, yeast tRNA, dextran sulfate and RNAse inhibitor to make hybridization mix. The samples were hybridized with hybridization mix overnight in a humid chamber at 37°C. The samples were washed two times with 1X SSC buffer (sodium saline citrate) containing 10% formamide and a final wash was done using 1x TBS buffer (Tris-buffered saline). Samples were mounted using SlowFade™ Diamond Antifade Mountant (ThermoFisher Scientific, S36967).

Fluorescence was detected by spectral unmixing of autofluorescence spectra using laser scanning confocal microscopy on a Zeiss LSM 880 multiphoton confocal microscope. We used 15% 561 nm laser and alpha Plan-Apochromat 100x/1.46 oil DIC lens to acquire *TDF in situ* images using the online figure print mode. Pure TxRed was used as positive control, and autofluorescence from asparagus tissue was used as negative control for spectral bleed-through. We used the 561 nm laser to detect *SOFF* mRNA labeled with TMR-linked probes, with the alpha Plan-Apochromatc 100x/1.46 Oil DIC lens, but in spectra mode. After images were taken, each image was spectra unmixed. Spectral data for the pure TMR fluorophore were used as positive control, and non-labeled samples were used as negative control spectra for spectral bleed-through. The brightness and contrast of images in the same figure panel were adjusted equally and linearly in Zen 2010 (Carl Zeiss).

### Image quantification

For quantification, the number for each localization events were calculated using Volocity (Perkin Elmer). For each image, we first removed noise using the default value from Volocity, then quantified the localization events using a pipeline consist of Find Object (in the channel with *in situ* signal), Separate Touching Objects (0.05 μm^3^), Exclude object (<0.05 μm^3^), and Exclude Object (>0.8 μm^3^). Five replicates for each z-stack of images were used for calculating the copy number for each gene. P-value was calculated using a t-test assuming equal variance.

---

**Supplemental Figure 1.**
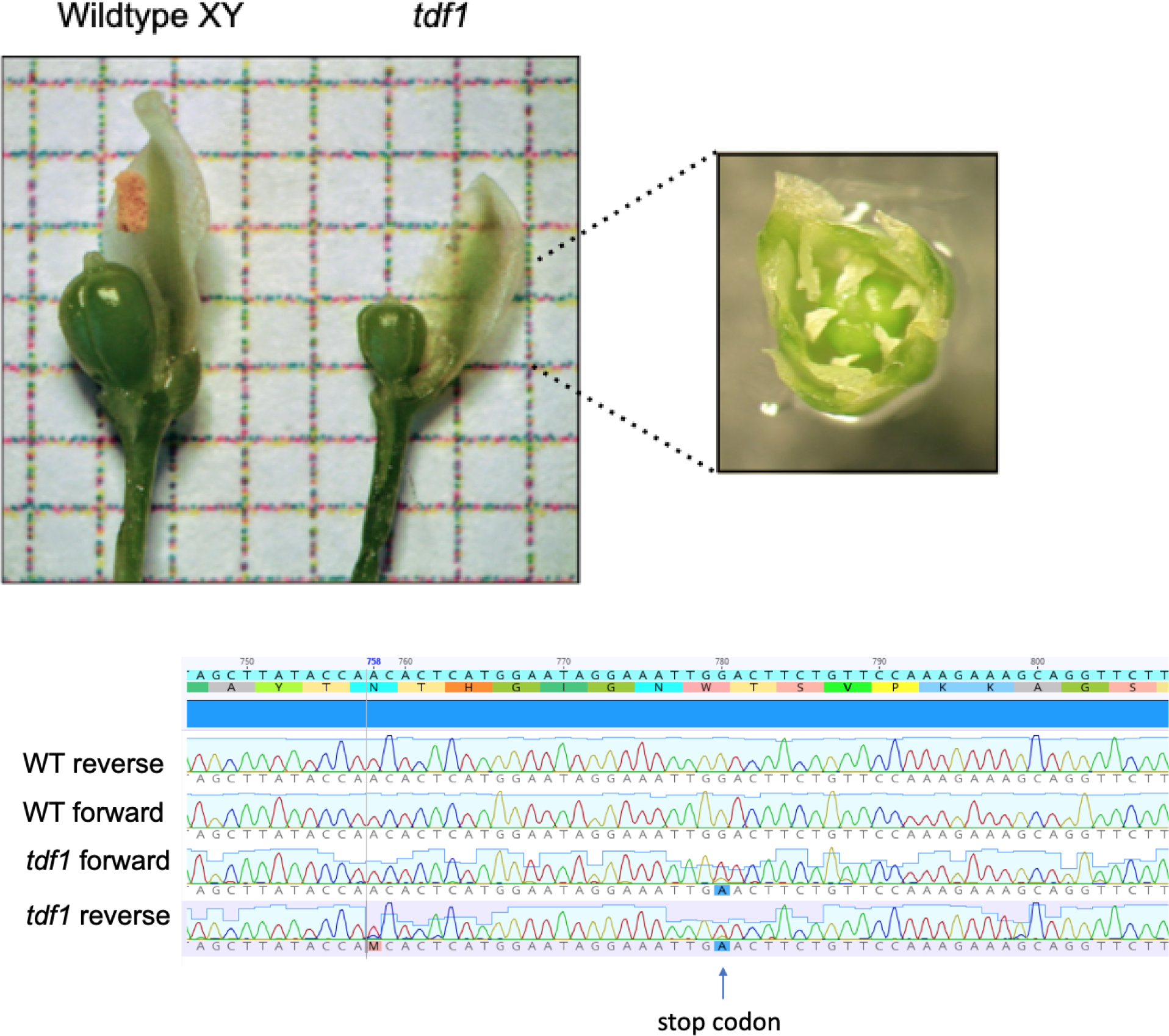
A) The EMS-induced *tdf1* mutant lacks anthers and is functionally neuter. B) Sanger resequencing of the wild type and *tdf1* mutants, highlighting the EMS-induced stop codon in the TDF1 gene.

**Supplemental Figure 2:**
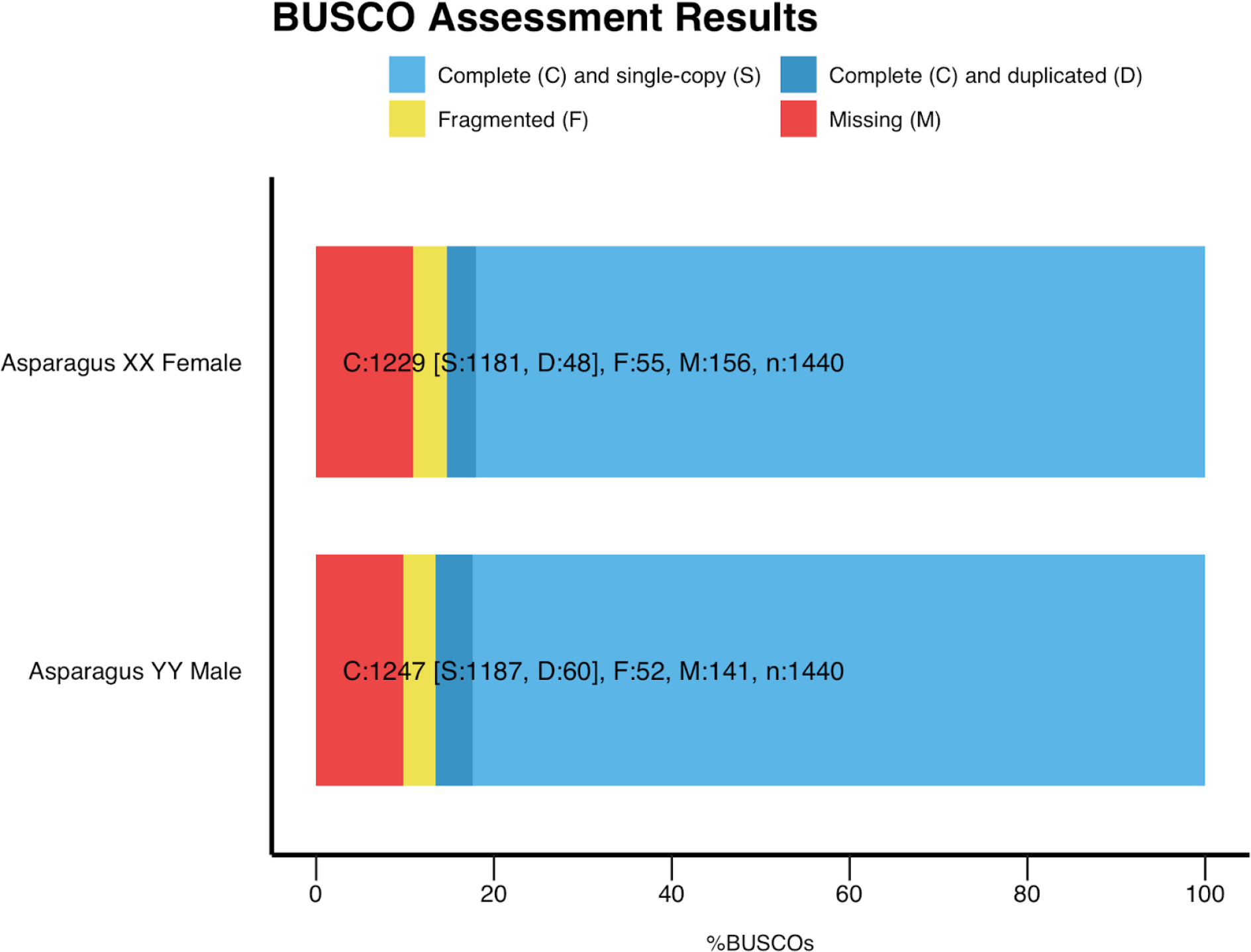
BUSCO score comparison for the previously published YY genome (*6*) and this manuscript’s XX genome assemblies.

**Supplemental Figure 3:**
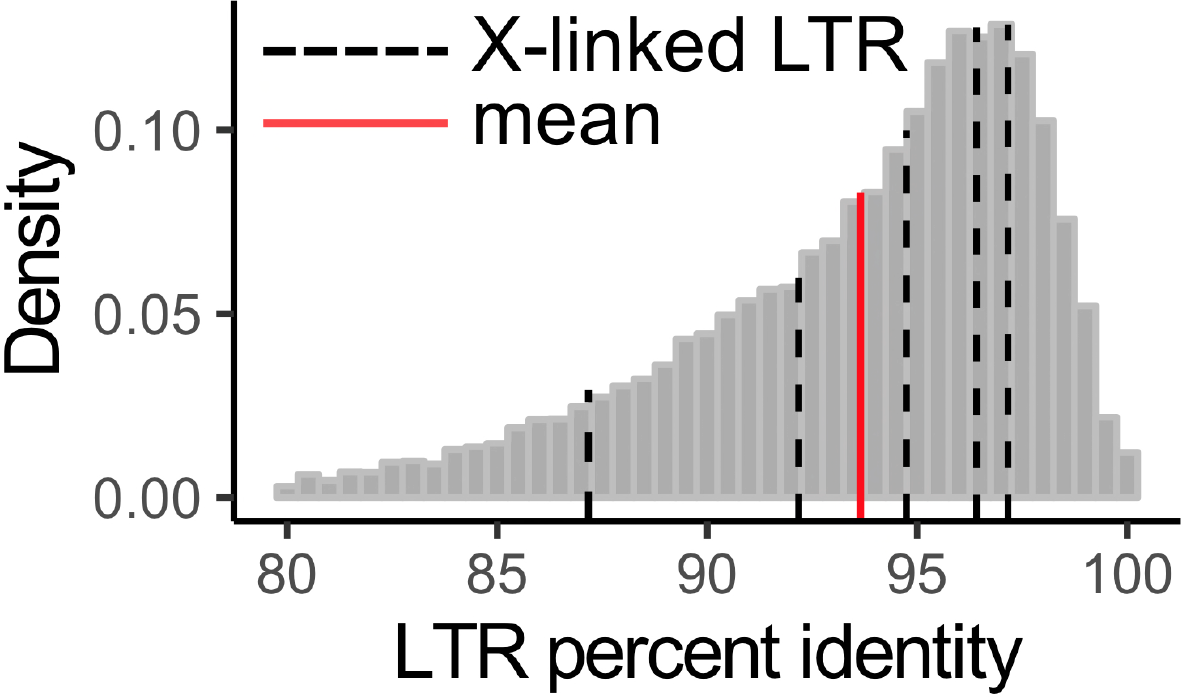
Distribution of X-linked LTR retrotransposon pairwise LTR percent identity comparisons (black dotted lines) compared to the genome-wide distribution (grey bars) and mean (red line).

**Supplemental Table 1:** Xlinked BLASTX (nr) gene annotations

| Gene ID | BLASTX annotation |
| --- | --- |
| MSTRG.231 | Zinc Finger protein WIP2 / NO TRANSMITTING TRACT (NTT) |
| MSTRG.232 | Uncharacterized protein |
| MSTRG.233 | Uncharacterized protein |
| MSTRG.234 | Outer envelope protein 80, chloroplastic |
| MSTRG.235 | Uncharacterized protein |

